# Serial Dependence in Temporal Perception Reveals the Dynamics of Constructing an Internal Reference Frame

**DOI:** 10.1101/2021.05.31.446431

**Authors:** Tianhe Wang, Yingrui Luo, Ernst Pöppel, Yan Bao

## Abstract

Temporal perception is crucial to cognitive functions. To better estimate temporal durations, the observers need to construct an internal reference frame based on past experience and apply it to guide future perception. However, how this internal reference frame is constructed remains largely unclear. Here we showed the dynamics of the internal reference construction from the perspective of serial dependence in temporal reproduction tasks. We found the current duration estimation is biased towards both perceived and reproduced durations in previous trials. Moreover, this effect is regulated by the variability of sample durations. The influence of previous trials was stronger when the observers were exposed to context with more variable durations, which is inconsistent with previous theories that the similarity between successive stimuli induces serial dependence. We proposed a Bayesian model with an adaptive reference updated continuously after each observation, which can better explain the serial dependence observed in temporal perception.

## Introduction

Temporal processing is essential to cognitive functions such as anticipation of future events or movement control^1–5^. Behavioral studies indicate that the perception of the temporal durations can be strongly influenced by past experience. One and a half century ago, with a series of temporal reproduction experiments, Vierordt found that the reproduced duration had a tendency to regress to the “mean” of all the sample durations presented in one experiment^6,7^. This phenomenon later referred to as Vierordt’s Law was proved to be universal in different perceptual modalities^8–10^ and various ranges of durations from hundreds of milliseconds to tens of seconds^11,12^. The recent theories to explain Vierordt’s Law proposed that the observers construct an internal reference based on the context durations they previously experienced^13–16^. The estimation of the current duration is an integration of current perceptual input and the internal reference conveying the contextual information^14,16–18^. This integration biases the estimation towards the mean of the internal reference, resulting in the phenomenon of regression-to-mean in the reproduced durations. However, it remains unclear how the internal reference in temporal perception is constructed and updated.

To examine the dynamics of the internal reference, a suitable way is to focus on how the trial history influences the current performance^15,19^. Specifically, different sample durations in a 1-back trial may change the internal reference and influence perception and reproduction of the current duration. The idea of a local contextual effect known as serial dependence has been examined in the visual domain^20^. For example, in a visual orientation judgment task, the perceptual reports of the current orientation are attracted toward the stimulus of the recent past^20–23^. Similar effects are observed in other perception tasks^21,23–26^ like face perception^27,28^ and numerosity perception^29,30^. In the temporal domain, the sequential effect has been examined in temporal judgment that a duration will be more likely judged shorter if it is preceded by a short duration^31,32^. A follow-up study showed that reproduction to the current stimuli would be slightly longer when participants reproduced to a 1200 ms stimuli than to a 300 ms stimuli in the previous trial^15^. These observations further motivated us to investigate serial dependence in temporal perception systematically using a temporal reproduction with a set of stimuli randomly sampled from a certain distribution. Indeed, in our temporal reproduction task, we found a nonlinear serial dependence whose pattern is similar to the serial dependence in the visual tasks^23-26^. Specifically, the current reproduction is biased towards the duration of the previous stimulus, and the bias first increased as the difference between stimuli increased but then dropped down when the difference was too large. By two follow-up experiments isolating the influence of motor reproduction and perception, we found that motor reproduction contributes mainly to serial dependence, while perception also has a moderate effect but was much weaker.

Besides the internal reference model, an alternative model proposed by previous studies was that serial dependence happens because participants believe two successive stimuli are generated from a common source and thus integrates them when making the current perception^7,33,39^. This integration model has been successfully applied to explain serial dependence in numerosity perception^39^. This integration model could somehow be regarded as an extreme case of the internal reference model whose reference is simply the stimulus duration in the 1-back trial other than a reference constructed based on whole trial history. Therefore, the key difference between the integration model and the internal reference model is whether the serial dependence would be influenced by the overall distribution of the stimulus besides the one back trial.

To compare the internal reference model and the integration model, we manipulated the range and the variability of the stimulus distribution and examined whether those changes influence serial dependence in the temporal reproduction task. Indeed, we found that the curve of serial dependence is strongly impacted by the stimulus context. A stronger serial dependence was found under a context with a wider stimulus range or a higher stimulus variability, which is more consistent with the internal reference model. Based on our behavioral results, we propose a Bayesian Recency model that suggests participants apply an adaptable internal reference updated towards the previous stimuli to facilitate the processing of the current duration. By model simulations, we show that our model can well explain different properties of serial dependence observed in the behavioral results.

## Result

### Temporal perception is biased towards the previous stimulus

To investigate how the recent trial history influences temporal perception, we examined the serial dependence in a ready-set-go temporal reproduction task^16^. Participants perceived the interval between two stimuli (“Ready” and “Set”) and pressed a button to make reproduction (“Go”; Fig. 1a). The sample durations were randomly sampled from a uniform distribution from 500 to 900 ms. Similar to most previous studies in temporal perception^14,15,31^, the participants’ reproduced duration showed a strong regression-to-mean, that they over-reproduced the short sample durations and under-reproduced the long sample durations (Fig. 2a). The slope of the reproduced duration to the sample duration was significantly smaller than one (0.65 ± 0.20, t(11) = -6.08, p<0.001). To measure potential serial dependence, we applied an index called deviation: the difference between the present reproduction and the individual’s averaged reproduction of the current sample duration (see Fig. 2a). This index can rule out the effect of regression-to-mean and overall reproduction bias from later analyses^32,33^.

**Figure 1.**
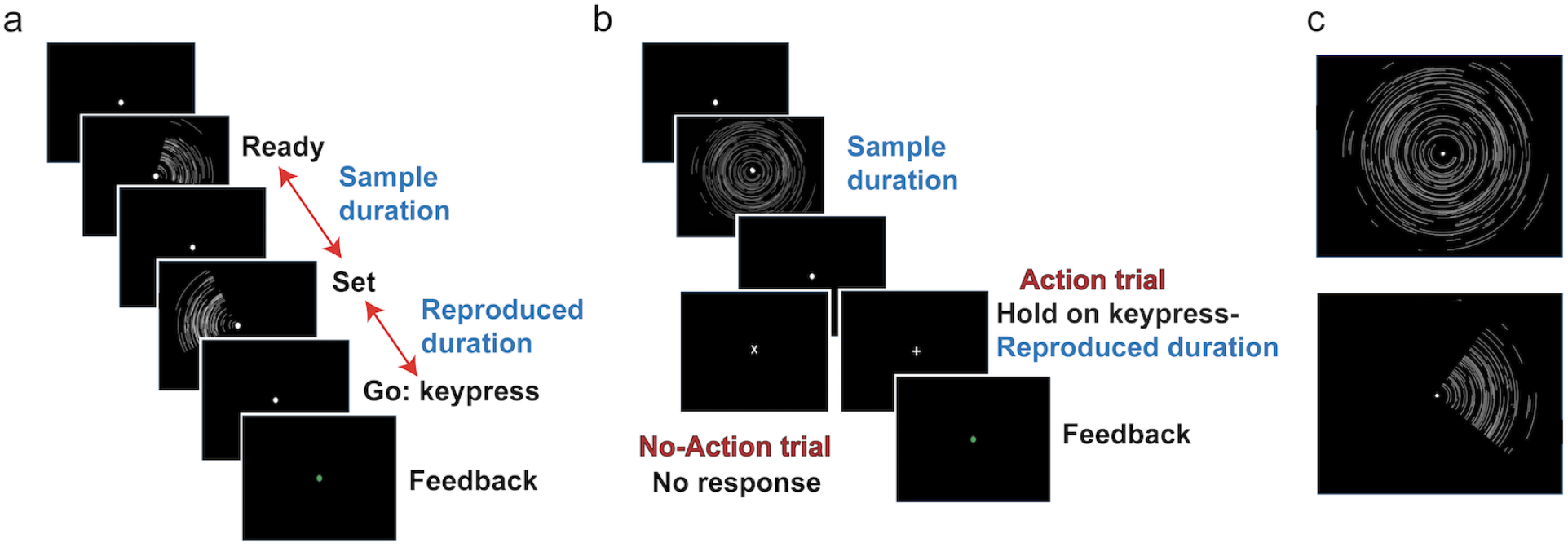
Event sequence and stimuli. (a) The stimulus sequence in Experiment 1. Each trial started with a fixation point at the center of the screen. The sample duration was presented as two 100-ms fan-shaped stimuli separated by the sample duration, with the first stimulus signifying “Ready” and the second stimulus signifying “Set”. Participants were instructed to press a key when they felt the interval between the onset of “Set” and the keypress the same as the sample duration. The reproduced duration was the interval between the onset of the “Set” signal and the keypress. To encourage better performance, the feedback was given at the fixation point after the keypress, with green as relatively accurate, red as inaccurate. (b) The stimulus sequence of Experiments 2-5. A ripple-shaped stimulus was presented for sample duration. After it disappears, the fixation point changed shape, with “+” indicating that participants should hold a keypress to reproduce the stimulus (action trial) and “x” indicating not to make a response (no-action trial). Participants hold the keypress and released it when they feel the pressing time (reproduced duration) the same as the sample duration. After the key was released, the fixation point changed color to give feedback. In the no-action trial, a new trial started after presenting the “x” for 700 ms. (c) A sample of stimulus in Experiments 2 - 5 (up) and Experiment 1 (down). The stimulus in Experiment 1 was a random quarter of the stimulus in Experiment 2-5. The “Ready” and “Set” signals in one trial did not share a common area in Experiment 1.

**Figure 2.**
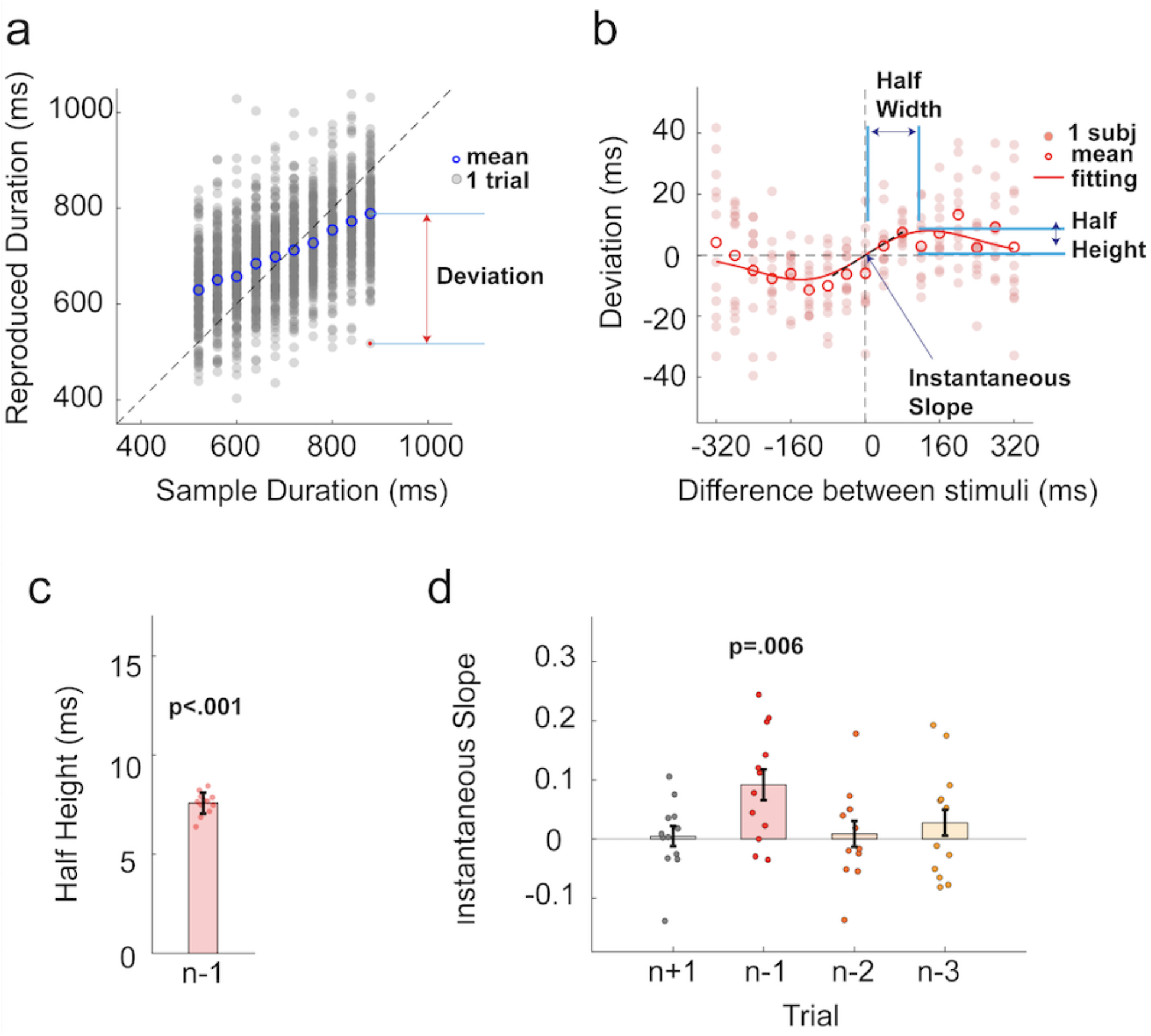
Results of Experiment 1. (a) Reproduced durations of a typical participant. Each grey dot represents the reproduced duration of one trial. The blue circle represents the averaged reproduced duration of a sample duration condition. (b) The nonlinear relationship between deviation (current reproduced duration minus the averaged reproduced duration) and difference (current sample duration minus precious sample duration) between stimuli. Each filled dot represents the averaged deviation of a participant. Each blank circle represents the average of 12 participants. The red line represents the best-fitted DoG when 12 participants are collapsed. (c) Half-height of the best-fitted DoG for the 1-back trial. Each filled dot represents an estimation in the jackknife resampling. Error bars represent standard error. (d) Instantaneous slope of DoG-fitting curve for 1-back (n-1), 2-back (n-2), 3-back (n-3), and 1-future(n+1) trials. Each filled dot represents the instantaneous slope of one participant. Bars indicate the average, and error bars represent standard error. The p-value represents a significant difference from zero.

In the present experiment, we saw a positive and nonlinear serial dependence between the sample duration difference and deviation (see Fig. 2b). That meant the present reproduced duration was more likely to be longer than the average when the previous sample duration was longer than the present one, and was more likely to be shorter when the previous sample duration was shorter than the present one. The magnitude of this effect was first raised when the difference between the sample durations increased but it then decreases when the difference became too large. Then we fitted serial dependence at the group level with a derivate of Gaussian (DoG), which was proved to be a good model describing the non-linearity of serial dependence^20,32^. The best-fitted DoG had a positive half-height of 7.57 ± 0.56 ms (Fig. 2c, t(11) = 13.4, p<0.001), quantifying the magnitude of the serial dependence. The DoG showed a better fitting of the group-level serial dependence than the non-model (y = 0, ΔAIC_n_ = -46.3 ± 7.7) and the linear model (y = kx, ΔAIC_l_ = -18.6 ± 4.6), indicating serial dependence in temporal reproduction to be indeed a nonlinear effect. We then examined how many previous trials serial dependence can transmit by fitting DoG to individual participants. A positive instantaneous slope (0.092 ± 0.094, t(11) = 3.39, p=0.006; Fig. 2d) was found in the 1-back trial, but no significant slope was found in 2-back (0.009 ± 0.079, t(11) = 0.39, p=0.70) or 3-back (0.028 ± 0.095, t(11) = 1.00, p=0.34) trials. Thus, a significant serial dependence could only be seen for the 1-back trial in Experiment 1.

### Serial dependence is mainly contributed by the motor response

Experiment 1 showed a nonlinear attractive serial dependence in the temporal reproduction task. In Experiment 2, we further asked whether this serial dependence was caused by the perception of the sample duration or the motor reproduction^28,32,34,35^. To dissociate the effects of perception and reproduction, we included 40% of no-action trials, where participants passively observed the sample duration without making a reproduction. By focusing on the action trial immediately followed a no-action trial, we could analyze the influence of the previous perception without the interference of the reproduction. Since the techniques concerns that an additional cue informing participants whether to make a response to the stimulus is easy to be confused with the “Ready” / “Set” signals and interrupts the reproduction, in Experiment 2, the sample duration was given by the presentation time of a static stimulus (see Fig. 1b-c), and reproduction was made by holding a keypress, i.e., the time between the keypress and the key-release is recorded as the reproduced duration.

A positive serial dependence was found when the 1-back trial was an action trial (instantaneous slope: 0.114 ± 0.042; Wilcoxon test: z = 2.03, p = 0.021; Fig. 3a). However, no serial dependence was found when the 1-back trial was a no-action trial (instantaneous slope: 0.006 ± 0.049; Wilcoxon test: z = 0.01, p = 0.97; Fig. 3a). Thus, the attractive serial dependence in Experiment 1 was presumably contributed mainly by the reproduction. The passive observation did not result in an observable serial dependence in the present task. Furthermore, when the 1-back was a no-action trial, a positive instantaneous slope was found (0.080 ± 0.034, Wilcoxon test: z = 2.25, p=0.012; Fig. 3a) if the 2-back trial was an action trial, which confirms the robust effect of reproduction. DoG fitting showed the half-height (7.42 ± 0.91 ms; Fig. 3b) and the half-width of the 1-back action trial (111.0 ± 15.6 ms) were very close to those in Experiment 1, although the task design was slightly changed. The 2-back action trial showed a smaller half-height (3.14 ± 0.60 ms) than the 1-back action trial (t(11) = 5.29, p =< 0.001; Fig. 3b), suggesting that the 1-back no-action trial weakens the attraction effect of the 2-back action trial.

**Figure 3.**
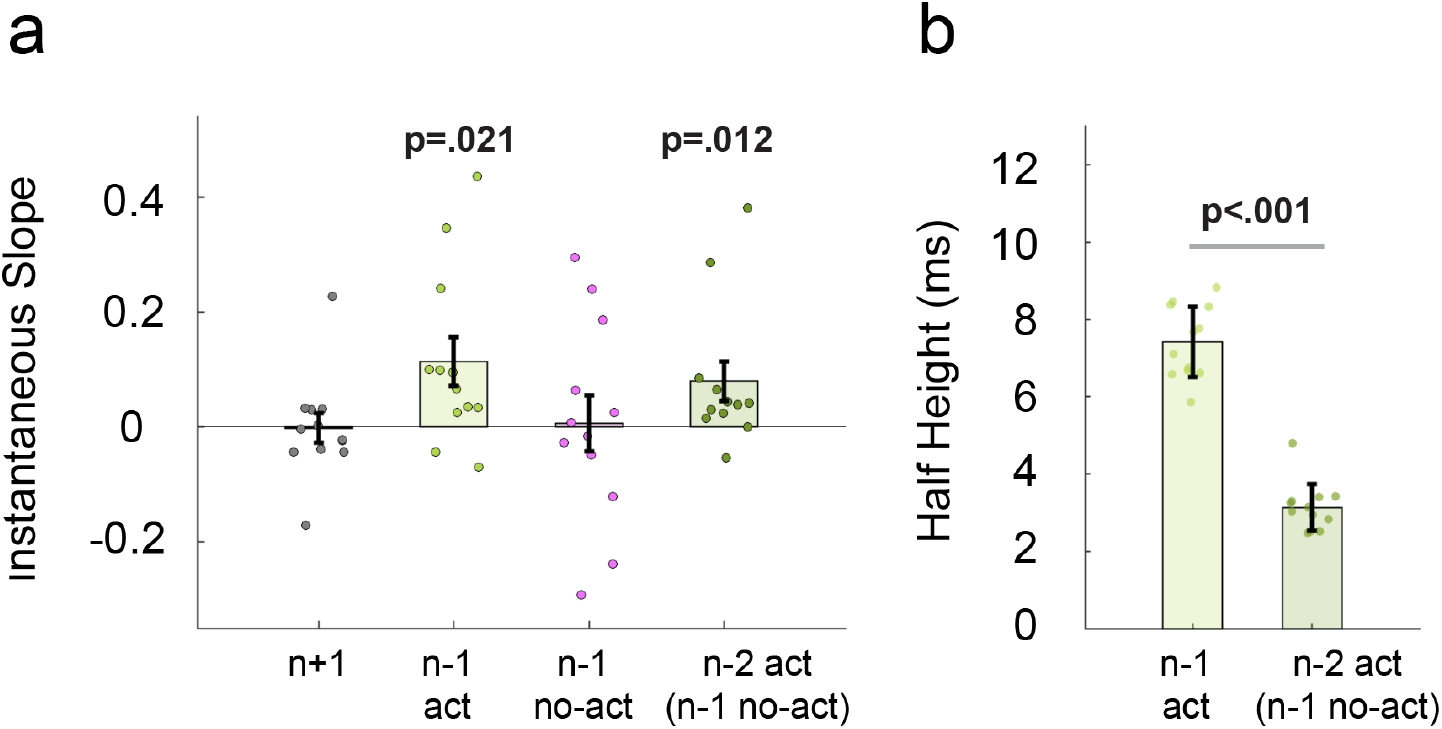
Results of Experiment 2. (a) Instantaneous slope of DoG-fitting curve for 1-back (n-1), 2-back (n-2), and 1-future (n+1) trials. Each filled dot represents the instantaneous slope of one participant. Bars indicate the average, and error bars represent one standard error. The p-value shows a significant difference from zero. Act: action trial; no-act: no-action trial. (b) Half-height of the best-fitted DoG for 1-back and 2-back trials. Each filled dot represents an estimation in the jackknife resampling. Error bars represent standard error.

### Perception without motor response has a weak attractive effect

It is necessary, however, to mention that participants were exposed to the duration twice in the action trial (perception and reproduction), but only once (perception) in the no-action trial. This difference might cause different results in serial dependence analyses of action and no-action trials. Moreover, serial dependence originating from perception was possibly not captured in Experiment 2 perhaps due to the insensitive measurement we used. Thus, to amplify the potential serial dependence of perception, we increased the percentage of no-action trials in Experiment 3 (see Fig. 4a). Sample durations in the action trials were specified as 700ms and preceded by 0-5 no-action trials either all longer (long prior condition) or all shorter (short prior condition) than 700ms. We confirmed that no participants realized that the sample duration in the action trial was fixed from the post-experiment inquiry. When preceded by 1-4 no-action trials, no significant difference between reproduced durations was found between long and short prior conditions (1-3 no-action trials: t(11) = 0.76, p = 0.46; 4 no-action trials: t(11) = 0.14, p = 0.89; Fig. 4b). However, the reproduced duration in the long prior condition was significantly longer than that in the short prior condition when preceded by five no-action trials (difference: 16.14 ± 18.20 ms, t(11) = 3.07, p = 0.011; Fig. 4b). These results suggested that the perceptions in the no-action trials weakly attracted the reproduction in the action trial, and it can only be observed when the effect has been accumulated, such as five no-action trials preceded before the current action trial.

**Figure 4.**
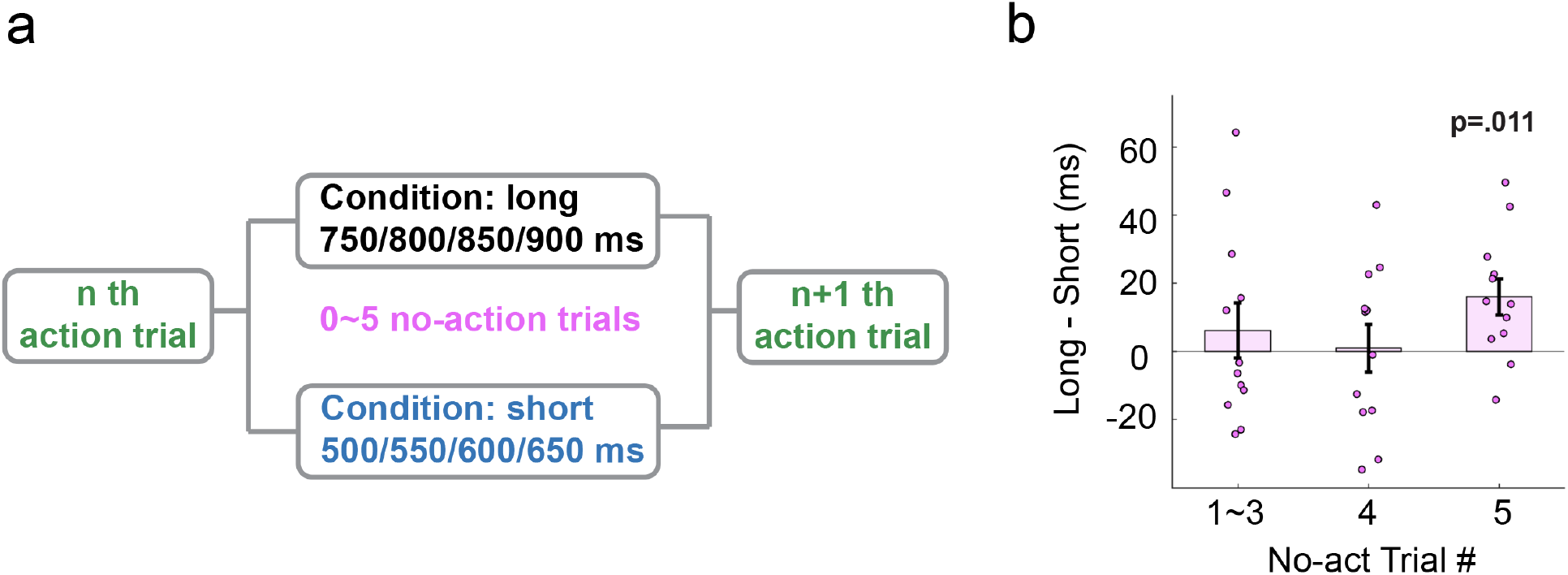
Design and result of Experiment 3. (a) Design of Experiment 3. Zero to five no-action trials either all longer (750-900ms) or all shorter (500-650ms) than 700ms was inserted between every two action trials. (b) The difference of reproduced durations between the long and the short prior conditions across different numbers of no-action trials preceded the current action trial. The p-value represents a significant difference from zero.

### Extending the range of stimuli enhances serial dependence in temporal perception

The serial dependence observed in Experiment 1-3 suggested that participants might update an internal reference to the reproduced duration and perceived duration after each trial. The perception in the next trial will be biased towards the new internal reference. However, an alternative hypothesis of serial dependence is that participants simply integrated the duration of the current and the 1-back trial when they think the two successive stimuli are generated from a common source^8,30,36^. The probability to perform the integration reduces as the difference between stimuli increases. Since this integration model supposes participants only use information from the 1-back trial to make the current perception, it predicts the serial dependence to be independent with the overall distribution of the stimuli as long as the difference between the previous and current stimulus remains the same. This prediction is different from the prediction of the internal reference model that serial dependence might be strongly influenced by the stimulus distribution.

To compared between the internal reference model and the integration model, in Experiment 4, we performed two temporal reproduction tasks with wider stimulus ranges (median-range: 540-1260 ms; long-range: 560-1840 ms) than that in Experiments 1&2 (short-range: 520-880 ms). Inconstant with the prediction of integration model, we found the amplitude of serial dependence increased as the range of stimulus extended. The best-fitted DoG of the median-range (ΔAIC_n_=-27.5 ± 4.2; ΔAIC_l_ = -10.8 ± 2.2) had a higher and wider DoG-shape curve than the short stimulus range (Fig. 5a). The best-fitted DoG of the long-range (ΔAIC_n_ = -20.7 ± 4.4; ΔAIC_l_ = -10.0 ± 2.1) showed a widest and highest DoG curve among three conditions (Fig. 5b). The half-height increases consistently (Fig. 5c; median-range: 11.8 ± 0.9 ms; long-range: 22.6 ± 2.0 ms; ts > 4.9, ps < 0.001). Similarly, the half-widths also increased constantly (ts > 10.5, ps < 0.001), indicating the current perception could be influenced by a more distant 1-back duration in conditions with a wider stimulus range. Moreover, the instantaneous slope of individual participants shows a significant positive serial dependence in not only the 1-back trial but also the 2-back trial for both the median-range (Fig. 5d, 1-back: 0.072 ± 0.094, Wilcoxon test: z = 2.60, p = 0.005; 2-back: 0.058 ± 0.092, Wilcoxon test: z = 2.21, p = 0.013) and long-range (Fig. 5e, 1-back: 0.093 ± 0.119, Wilcoxon test: z = 2.31, p = 0.010; 2-back: 0.070 ± 0.076, Wilcoxon test: z = 2.60, p = 0.005), which further indicated that sequential effect became stronger in wider stimulus range.

**Figure 5.**
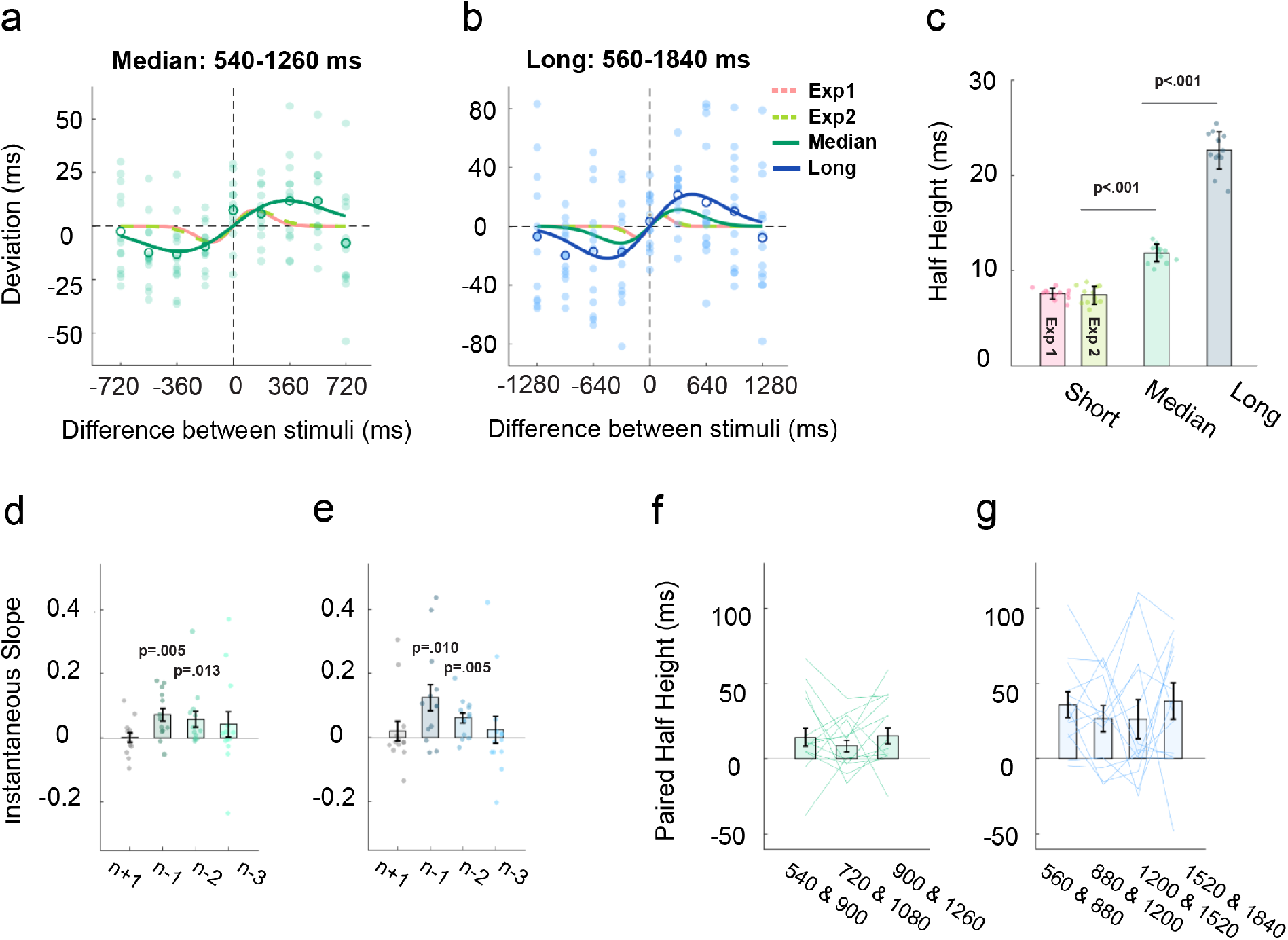
Results of Experiment 4. (a) Serial dependence of the 1-back trial in median range condition. Each filled dot represents the averaged deviation of a participant. Each blank circle represents the average of all participants. The turquoise line represents the best-fitted DoG when 13 participants’ data are collapsed in the median range condition. The red and green dash lines represent the best-fitted DoG of the n-1 trial in Experiment 1 and 2, respectively. (b) Serial dependence curves of the 1-back trial of long-range condition. The blue line represents the best-fitted DoG when 13 participants’ data are collapsed in the long-range condition. (c) Half-height of the best-fitted DoG for the n-1 trial in Experiments 4, 1 & 2. The pink bar represents Experiment 1. The green bar represents Experiment 2. Each filled dot represents an estimation in the jackknife resampling. Error bars represent standard error. (d-e) Instantaneous slope of DoG-fitting curve for 1-back (n-1), 2-back (n-2), 3-back (n-3), and 1-future (n+1) trials in median (d) and long (e) range conditions. Each filled dot represents the instantaneous slope of one participant. Bars indicate the average, and error bars represent one standard error. The p-value shows a significant difference from zero. (f-g) Serial dependence of each stimuli pair in median (f) and long (g) range conditions. Each line represents one participant. Bars indicate the average, and error bars represent one standard error.

A plausible explanation for the increase of serial dependence in the wider stimulus range is the scalar property in time perception. That is, participants may have larger perceptual noise for a longer duration and in turn, tend to rely more on the previous response when perceiving the long standard duration. If this explanation holds true, the serial dependence would be stronger as the magnitude of the current stimulus increases (fig. 6k upper panel). To test this hypothesis, we selected a series of stimuli pairs with the same distance but different averaged durations, and calculated serial dependence of each pair (see method). However, we did not find a continuous increase in serial dependence across different stimulus pairs when the mean duration increased in the median-range (Fig. 5f, ANOVA: F(2, 24) = 0.76, p = 0.48) or long-range (Fig. 5g, ANOVA: F(3, 36) = 0.43, p = 0.73) conditions. This suggested that the increase of the serial dependence in a wider stimulus range cannot be fully explained by the increase of perceptual noise for the longer standard durations.

**Figure 6.**
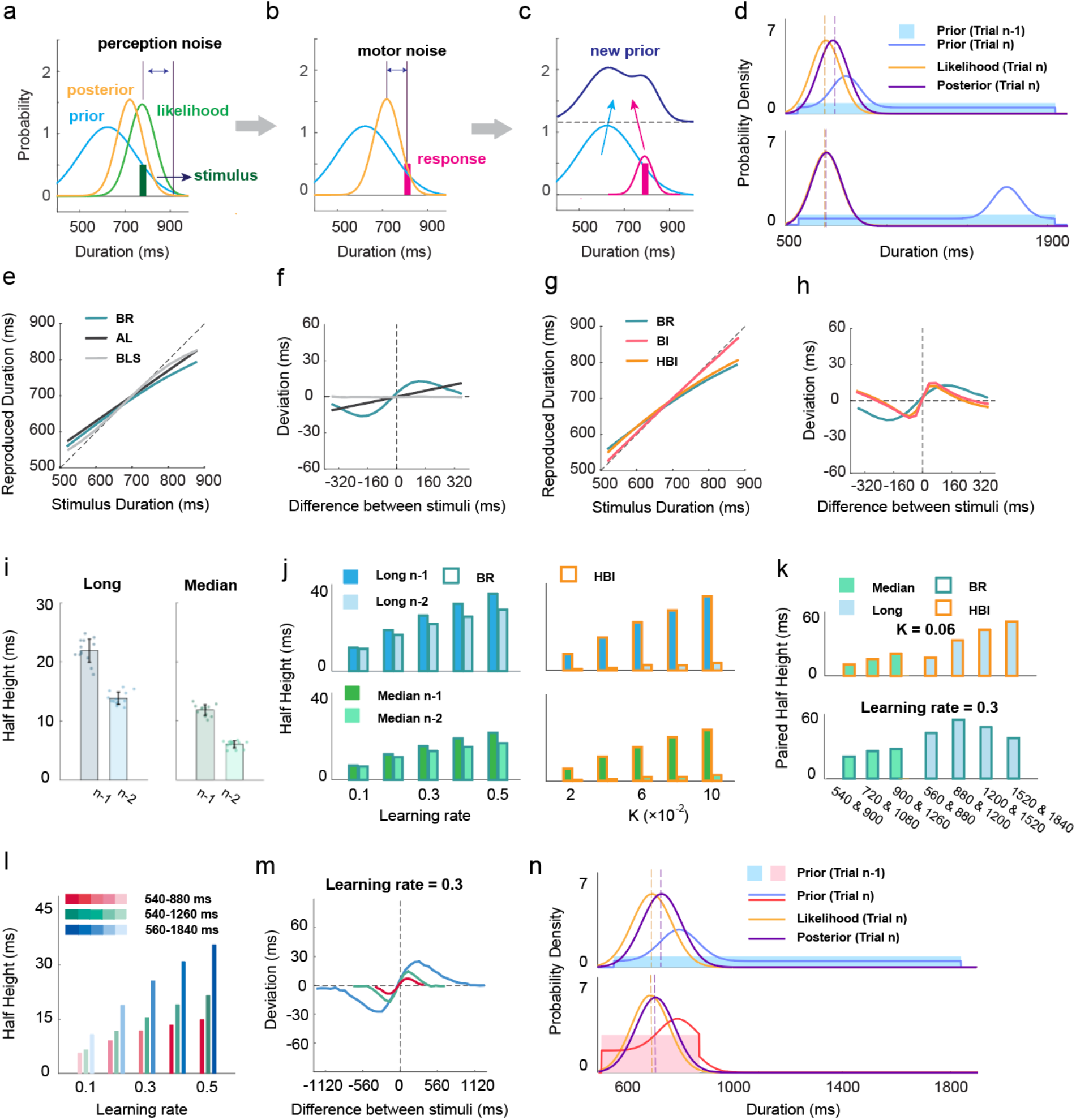
Model predictions (a-c) Illustration of the Bayesian Recency model. (a) The stimulus was transferred into a normal distribution (likelihood) centered at the sample duration with a variance of perception noise. The prior is multiplied with the likelihood to estimate the posterior following the Bayesian rule. (b) The mean of the posterior was used as the final estimation of the sample duration. The reproduced duration followed a normal distribution centered at the estimation with a variance of motor noise. (c) After making the motor response, the observer updates the prior based on the reproduced duration by adding up a scaled normal distribution centered at the reproduced duration. (d) Illustration of how the BR model generates a non-linear serial dependence. Upper panel: the previous stimulus duration influences the prior around the current likelihood when they are close. Lower panel: the previous stimulus duration influences the prior far from current likelihood and has little contribution to the posterior. (e-f) The central tendency (e) and serial dependence (f) of the simulation results of the BLS, the AL, and the BR. (g-h) The central tendency (g) and serial dependence (h) of the simulation results of the BI, the HBI, and the BR. (i) Half-height of the best-fitted DoG for the n-1 and n-2 trial in long-range (left) and median-range (right) conditions in Experiments 4. Each filled dot represents an estimation in the jackknife resampling. Error bars represent standard error. (j) Simulation of (i) with the BR (left) and the HBI (right). The upper two panels are the simulation to long condition and the lower two panels are the simulation to median condition. (k) Serial dependence of stimuli pairs generated by HBI (upper) and BR (lower). (l-m) Simulation of BR predicting that the half-height of serial dependence increases as the range of stimulus extends. (n) Illustration of how the BR generates the prediction that the long stimulus range (upper panel) induced a stronger serial dependence than the short stimulus range (lower panel). The stimulus in the n-1 trial is supposed to be 800 ms. The learning rate is set as 0.7.

### Nonlinear serial dependence can be best explained by a Bayesian Recency model

We propose a computational model to explain the regression-to-mean and nonlinear serial dependence in the temporal reproduction task observed in our experiments. We hypothesize that participants hold an internal representation of the stimulus distribution and update this internal reference when a new sample is observed. To estimate the duration of the current stimulus, the perceptual input is integrated within the internal reference flowing the Bayesian rule (Fig. 6a-b). Since our Bayesian model continuously updated the internal reference based on the most recent trials (Fig. 6c), we named it the Bayesian Recency (BR) model. As for control, for model simulation, we also included a classic Bayesian model (Bayesian-last-square, BLS)^9^, which assumes the true distribution was learned as an internal reference and will not be updated during the experiment. Besides Bayesian models, we also included an adaptable model with a simpler format (AL model) based on Adaptation level theory^37^. AL model assumes the internal reference is a single value instead of a distribution and the observers make use of it by a weighted average between the internal reference and the perceptual input.

The AL, the BR, and the BLS model were applied to simulate participants’ behaviors in Experiment 1. All three models showed a regression-to-mean to some degree (Fig. 6e). Strikingly, the three models showed a large difference in predicting serial dependence (Fig. 6f). Both the AL and the BR models were able to predict an attractive serial dependence, while the BLS model did not generate any serial dependence. Importantly, only the BR model predicted a DOG-like nonlinear serial dependence, whose shape was very close to the results in human participants. In the BR model, when the current and previous durations are close, the previous duration can influence the shape of prior in the range around the current duration and make the current posterior shift more relative to likelihood (Fig. 6d upper panel). However, when the current and previous duration are distinct, the previous duration cannot influence the shape of the prior around the likelihood, so the attraction effect decreases (Fig. 6d lower panel). The AL model showed a linear serial dependence such that the deviation strictly increases as the difference between the durations increase. It is clear that the BR model can better explain the serial dependence than the AL and the BLS models, which supports the assumption that the observes hold a dynamical and distributional internal reference of durations rather than a fixed or a single-value reference.

Besides our BR model, we also implemented the Bayesian Integration (BI) model from the previous study, which supposes the observers integrate current perceptual input and the previous response^30^. The probability to integrate the previous response is determined by the difference between two successive durations that a smaller difference means a higher probability of integration. This model also predicts a non-linear serial dependence (Fig. 6h). However, when controlling the size of serial dependence to a reasonable range, the BI model could not show a regression-to-mean (Fig. 6g). Thus, we further built up a Hierarchical Bayesian Integration (HBI) model, which supposes the observers perform the first Bayesian integration between the current perceptual input and the internal reference to get an estimation, and then perform a secondary Bayesian integration between this estimation and the previous response. The HBI model predicts a similar central tendency pattern as the BR with the first Bayesian integration and a serial dependence like BI with the second Bayesian integration. Note, similar to the BLS, the HBI model also assumes a fixed internal reference. One critical difference between the BR and the HBI (or BI) is that the HBI (BI) model cannot predict a serial dependence appropriately before the 1-back trial since only the 1-back response is integrated (Fig. 6j right). However, in the BR model, serial dependence can pass on for a few trials by the internal reference (Fig. 6j left). We saw a considerable serial dependence from the 2-back trial in both median-range and long-range conditions (Fig. 6i), which is consistent with the prediction of the BR. This result supported the idea that serial dependence in temporal reproduction tasks is due to the update of the internal reference but not a simple integration between the two successive durations.

As for the influence of stimulus context, the BR model predicts that the half-height of serial dependence increases as the range of stimulus durations extends wider (Fig. 6l-m). In short-range, the observers construct a concentrated prior which is more resistant to the change and remains relatively flat after a new sample is added to the prior (Fig. 6n lower panel). Therefore, the posterior will not vary away too much from the likelihood after the Bayesian integration. On the contrary, the prior in long-range is more distributed and shows a more prominent local change after each updating (Fig. 6n upper panel). Therefore, the posterior will be influenced more by the previous trial, which results in stronger serial dependence. Although the HBI model also predicts the long condition has a stronger serial dependence than the median prior condition (Fig. 6j right), that effect is likely to be induced by the scalar property in time perception -- a larger perceptual noise for longer durations that results in a wider-range integration between the previous and the current durations. For the prediction of the HBI model, serial dependence consistently increases when the mean duration of the stimuli pairs increases (Fig. 6k upper panel). This is not consistent with the behavior results since the serial dependence does not increase dramatically across stimuli pairs (Fig. 5f-g). Consistent with the behavior result, we did not see a monotonous increase of serial dependence across stimuli pairs in the prediction of the BR (Fig. 6k lower panel).

### High contextual variability enhances serial dependence

To further confirm that it is the contextual variability that influences the serial dependence but not the perceptual noise in long durations, we added two more conditions to control either the mean duration of the context or the range of the stimulus durations based on the median-range condition in Experiment 4. We added a new short-range condition from 720 to 1180 ms (short-900 condition; Fig. 7a), which shared the same mean (900 ms) as the median-range (540 -1260 ms) and matched the width of the context in Experiment 1& 2 (360 ms). Additionally, we also added a bimodal-distribution condition with the same range as the median condition but had a higher contextual variability (Fig. 7a). The BR predicts stronger serial dependence in context with higher variability (Fig. 7b). That is, the half-height of the bimodal-distribution is larger than that of the median-range uniform prior, and the half-height of the median-range is larger than the short-900 with the same mean. HBI does not predict any difference between the median-range and the bimodal-distribution (Fig. 7c), and it predicts a moderate and inconsistent difference between the short-900 and the other two conditions. Table 1 summarized the predictions of different models on the properties of serial dependence we have mentioned so far.

**Figure 7.**
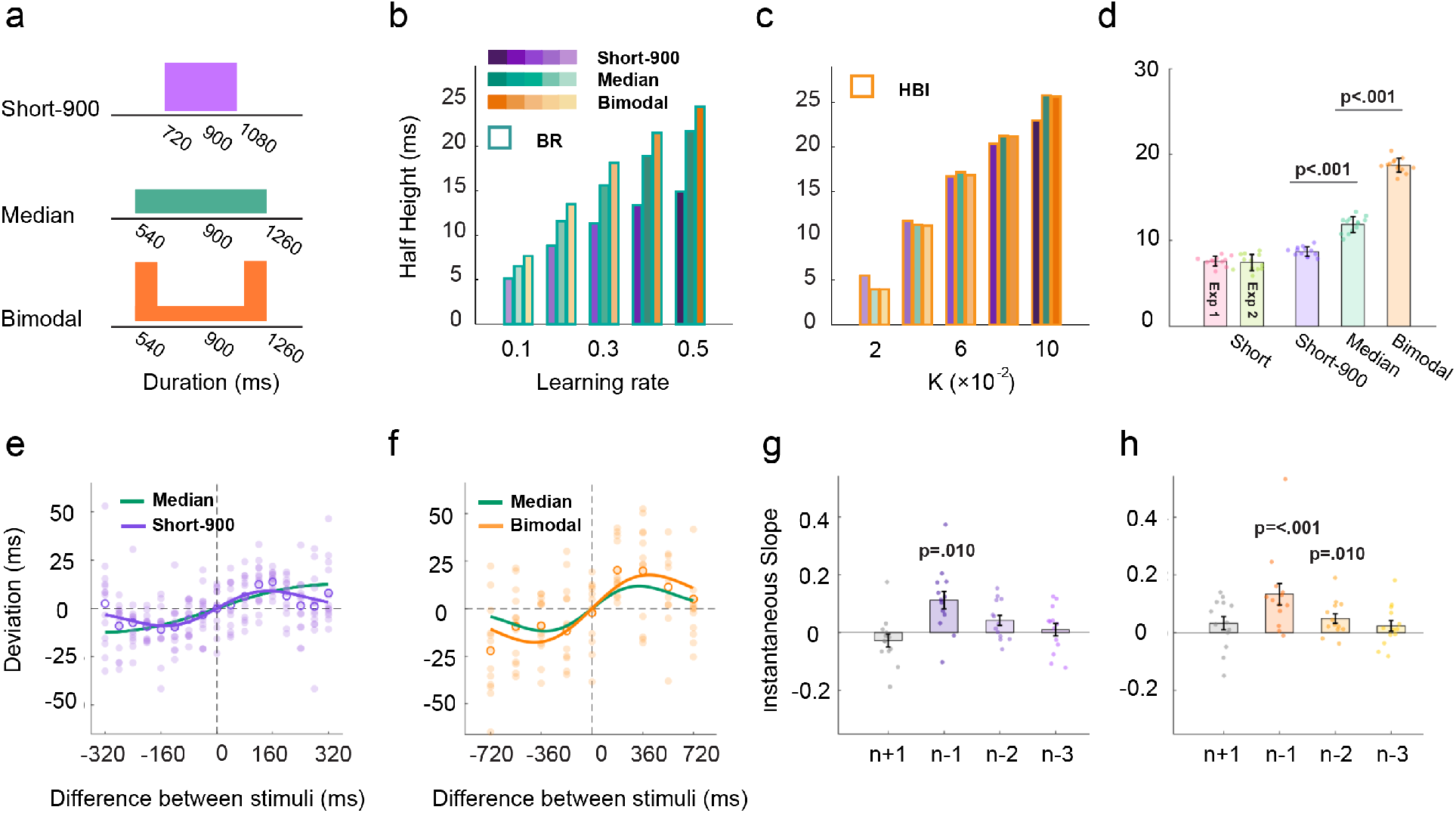
Results of Experiment 5. (a) Illustration of the distributions of stimulus durations in experiment 5. The short-900 condition (purple) has the same mean as the median-range (turquoise). The bimodal-distribution (orange) has the same range as the median prior but has a larger variability. (b-c) Half-heights of the best-fitted DoGs in different conditions predicted by the BR model (b) or the HBI model (c). (d) Half-height of the best-fitted DoG for the n-1 trial in Experiments 5, 1 & 2. Each filled dot represents an estimation in the jackknife resampling. Error bars represent standard error. (e) Serial dependence of the 1-back trial of the short-900 condition. (f) Serial dependence of the 1-back trial of bimodal distribution condition. Each filled dot represents the averaged deviation of a participant. Each blank circle represents the average of all participants. The purple line, orange line, and turquoise line represent the best-fitted DoG for the short-900, the bimodal-distribution, and the median-range uniform condition, respectively. (g-h) Instantaneous slope of DoG-fitting curve for 1-back (n-1), 2-back (n-2), 3-back (n-3) trials, and 1-future (n+1) trial in short-900 (g) and bimodal-distribution (h). Each filled dot represents the instantaneous slope of one participant. Bars indicate the average, and error bars represent one standard error. The p-value means a significant difference from zero.

**Table 1.** Predictions of different Models on serial dependence.

| Model/Prediction | Bayesian Recency (BR) | Adaptation Level (AL) | Bayesian-least-square (BLS) | Bayesian integration (BI) | Hierarchical Bayesian integration (HBI) |
| --- | --- | --- | --- | --- | --- |
| Regression-to-mean | Yes | Yes | Yes | Little effect | Yes |
| Shape of Serial dependence | DoG-like curve | Linear curve | No serial dependence | Non-linear curve | Non-linear curve |
| Influence from trial n-2 | Yes | Yes | - | Little effect | Little effect |
| How serial dependence changes when extending the context range while holding the average duration | Increase | Increase | - | Slightly increase or decrease | Slightly increase or decrease |
| How serial dependence changes when increasing the context variability while holding its range | Increase | No effect | - | No effect | No effect |

The results turned out to meet the predictions of the BR. We found a stronger serial dependence in the bimodal condition than in the median-range condition and the weakest serial dependence in the short-900. As for the best-fitted DoG curve (Fig. 7e-f; 720-880 ms: ΔAIC_n_ = -48.3 ± 7.8; ΔAIC_l_ = -21.0 ± 4.5; bimodal: ΔAIC_n_ = -78.6 ± 5.6; ΔAIC_l_ = -18.3 ± 3.6), the half-height of short-900 (8.7 ± 0.6 ms) was close to that of previous short-ranges, and importantly, was smaller than the half-height of the median-range (Fig. 7d, t(25)>4.2, p<0.001). Besides, the instantaneous slope in the short-900 condition only showed a significant positive serial dependence in the 1-back trial (0.113 ± 0.110, Wilcoxon test: z = 2.31, p = 0.010) but not in the 2-back trial (Fig. 7g, 0.043 ± 0.064, Wilcoxon test: z = 1.49, p = 0.068). Moreover, the bimodal-distribution showed a larger half-height (18.7 ± 0.8 ms) than the median-range (Fig. 7d, t(25) > 9.3, p<0.001). It showed significant serial dependences in both the 1-back (Fig. 7h, 0.121 ± 0.014, Wilcoxon test: z = 3.30, p<0.001) and 2-back trials (0.048 ± 0.016, Wilcoxon test: z = 2.31, p = 0.010). Thus, with both the mean and the range of the context being controlled, our results further support the idea that higher variability induces a larger serial dependence, which is consistent with the prediction of the BR model, but not the HBI model.

## Discussion

Temporal perception is strongly influenced by past experiences. A body of studies suggested that the observers build up an internal temporal reference frame from the stimulus context to guide future perception^13,16,38^. The present study investigated how this internal reference is constructed and updated from the perspective of serial dependence, i.e., how the recent trial history influences the current perception. We provided a systematic analysis of serial dependence in the temporal reproduction tasks with these results: 1) The current reproduction is biased towards the sample duration of the previous trial (Experiment 1); 2) This serial dependence is mainly determined by the motor response (Experiment 2), 3) Perception without a motor response causes a weak attractive serial dependence (Experiment 3). 4) The amplitude of serial dependence increases in a stimulus context with a higher variability (Experiments 4 & 5). 5) A Bayesian model with a trial-by-trial updated internal reference which we refer to as the Bayesian Recency model (BR) can better explain the features of nonlinear serial dependence observed in the behavioral results than alternative models (Bayesian-Least-Square, Adaptation-Level, Bayesian Integration, Hieratical Bayesian Integration). These results suggest that the estimation of the temporal duration is more likely to be based on a probability-based computation rather than a linear regression to a reference point. Furthermore, serial dependence is more likely to be a result of a continuous update of the internal reference rather than a simple integration between two successive trials.

Our results on serial dependence reveal two important features of the internal temporal reference: it is represented as a distribution and can be updated trial-by-trial. Both the Bayesian models^13,16,39,40^ and AL models^3,14^ can explain the phenomenon of regression-to-mean in temporal perception. The major difference between the Bayesian models and AL models is whether the observer performs a probability-based computation or a simple linear regression. The AL observer is simply biasing the perception to the reference point that reflects an internal representation of the “mean” value of the context. The Bayesian observer applies not only a “mean” value but also other information about stimulus distribution, such as the variance, which theoretically can help the observer to obtain a better estimation than the AL observer does. However, maintaining distribution and processing probability-based calculations can cause a higher calculational load compared to solely computing with a reference point. No previous study has tested which of those two frameworks can better explain temporal perception. Strikingly, Bayesian models and the AL model generate a clear different prediction of serial dependence in our experiments, which offers a good opportunity to compare them. The Bayesian Recency model can well predict the non-linearity of serial dependence in behavioral results, while a simple version of the AL model only generates a linear serial dependence. Admittedly, if we allow some parameters (e.g., learning rate) in AL models to change across trials based on the distance between the current stimulus duration and the reference point, it might generate some degrees of non-linearity, but it is still very difficult for this adapted AL model to produce a DoG-like curve as the behavioral data showed. Therefore, our results support the hypothesis that humans apply a probability-based computation, following a Recency-Bayesian model, to integrate the context information in temporal perception.

An important empirical result of our study is that we found serial dependence increased as the range or the variability of stimulus durations increased. We confirmed this result by changing the range of the stimulus durations with the mean controlled and changing variability with even the range controlled. This result contradicts some previous theories on central tendency and serial dependence^30,36^, which suppose the observers make use of the past experience by applying the 1-back trial as prior and integrate it with the current perceptual input to estimate the duration. We implemented that theory with the HBI (BI) model. The HBI model predicted the serial dependence looks very similar when the durations of the current and the 1-back trials remain the same regardless of the global stimulus duration context, which is inconsistent with the behavioral results. Our BR model with an adaptable internal reference can well explain the relationship between serial dependence and contextual variability. The BR model suggests that an internal reference with high variability is more distributed and therefore, less resistant to time. The local distribution alters substantially after updated with each new observation, thus resulting in a stronger serial dependence. Besides the contextual influence on serial dependence, our BR model can also predict the considerable serial dependence of the 2-back trial. Still, the BI and HBI models fail to predict this result.

One crucial question in serial dependence is what kind of perception or motor processes may cause the effect. Recent studies on visual perception found that serial dependence is presumably implemented as multiple processes^23,32–35^. A common result is that motor response induces an attractive serial dependence^20,23,33^, which is also the case in our temporal reproduction task shown in Experiment 2. This result is consistent with several previous studies showing motor reproduction or movement time of making a response can influence the temporal perception immediately afterwards^38,41–43^. Considering that neural timing systems share common movement-related brain regions like the cerebellum and the premotor cortex^5,18,44,45^, it can even be expected that the reproduction process has a determinable effect in the serial dependence of temporal perception.

However, previous studies on serial dependence showed inconsistent results when testing serial dependence in perception without motor response in visual perceptions^32,33^. It is even more difficult to predict this result under the context of the temporal perception since previous results on duration judgment implied opposite sequential effects. When participants are continuously exposed to one duration and suddenly perceive a new one, they tend to overestimate the difference between the old and the new durations, known as the adaptation effect, and predicting negative serial dependence^46–50^. However, studies in the duration judgment tasks show that the estimation of the current comparison duration is attracted by the one in the previous trial^51,52^. Moreover, inconsistent results were found in the recent studies regarding whether the passive observations have any influence on regression-to-mean of the duration reproduction in different studies^53,54^, which make it more confusing whether motor response is necessary to construct the internal reference frame. In Experiment 3, we indeed observed a significant attraction from the previous stimuli duration instead of an adaptation effect after amplifying the effect of perception by repeatedly presenting durations either all longer or all shorter than the target sample duration. This attraction effect confirms that the perception itself is sufficient to influence the internal reference^19,51,53^. However, the impact of perception is much weaker than the reproduction and can only be detected when repeated five times, which perhaps explained no impact of passive observation on central tendency in some previous study^54^. Besides, the no-action trials preceding the action trial were not a constant duration but could vary within a certain range in Experiment 3, which might explain the lack of adaptation effect in our results.

To conclude, we showed that the perception of current duration is biased towards previous durations in the temporal reproduction task. This serial dependence is implemented in both perception and motor response and is enhanced by the variability of stimulus duration context. The model simulation results provide evidence that the observer constructed a distributional representation of the internal reference that is updated continuously based on each new observation. Our results shed light on flexible mechanisms in the perceptual system of temporal perception, in particular, to take into account prior information from the environment.

## Method

### Participants

A total of seventy-eight undergraduate or graduate students at Peking University were recruited for five experiments. All participants were right-handed with normal or corrected-to-normal vision. In Experiment 1, thirteen participants (8 females, mean age = 21.1, SD ± 1.00) were recruited for the two-day experiment with an hour each day. One participant quitted the experiment. In Experiment 2, twelve participants (8 females, mean age = 24.1, SD ± 4.17) went through the two-day experiment with one and a half hours each day. In Experiment 3, fourteen participants (6 females, mean age = 21.7, SD ± 4.00) went through a one-hour experiment, and 2 participants were excluded for analyses because the equipment failed. In Experiment 4, twenty-six participants (15 females, mean age = 21.1, SD ± 2.14) were randomly assigned into median-range and long-range conditions (13 for each) and went through a one-hour experiment. In Experiment 5, twenty-six participants (19 females, mean age = 21.1, SD ± 2.02) were randomly assigned to the short-900 condition lasting for an hour or the bimodal-distribution condition lasting for two hours (13 for each). Each participant was paid 70¥/h for their time. All experimental protocols were approved by the ethical committee of the School of Psychological and Cognitive Sciences, Peking University, and carried out according to the approved guidelines. We obtained written informed consent from all subjects before their participation.

### General methods

In all experiments, participants viewed stimuli from a distance of 65 cm on a 27-inch LCD monitor at a resolution of 1,024 × 768 driven by windows 8 system computer at a refresh rate of 100 Hz in a dark, quiet room. The background of the screen was set as black. The stimulus of the experiment was either a quarter (Experiment 1) or a whole (Experiment 2-5) ripple-shaped pattern in grey that centered at the fixation point (see fig. 1C). Since continuously presenting the same stimulus will cause repetition suppression which makes the stimulus duration perceived to be shorter^55–59^, the pattern of the stimulus was changed after each trial, with the spatial density of brightness remaining the same among all stimuli within one experiment. The fixation point was a grey spot with a diameter of 0.5 degrees at the center of the screen. The task was controlled by a customized program written in MATLAB (Mathworks, Natick, MA; Psychophysics Toolbox).

One-sample t-tests and paired t-tests were applied at the group level for comparisons. Normality assumption and equal variance assumption were confirmed before t-tests. Wilcoxon Sign-rank test was applied when the normality assumption was violated. Two-tail P value was reported for all statistic tests, and the significance level was set as p < 0.05. All analyses were performed with MATLAB (The MathWorks, Natick, MA) and SPSS version 19 (IBM, Somers, NY).

### Experiment 1

#### Procedure and design

A ready-set-go time-reproduction task^16^ was applied in Experiment 1, where participants perceived the duration of an empty time interval presented by a pair of flashed stimuli and reproduced this duration by the keypress. Each trial started with a fixation point at the center of the screen. After a random interval ranged from 0.7-1.2 s (following an exponential distribution), two 100-ms stimuli flashed in sequence, signifying first “Ready” signal and then “Set”. Both stimuli were a quarter of a random-generated ripple-shaped pattern (see Fig. 1c) that changed trial by trial. Two stimuli appeared in random degrees but shared no area in common. The sample duration was the interval between the onset times of “Ready” and “Set” signals. Participants were instructed to perceive the sample duration and pressed the space bar when the interval between keypress and the onset time of the set signal was the same as the sample duration. After keypress, the fixation point changed color to present binary feedback to encourage better performance. Red indicated the reproduction was relatively imprecise and green indicated relatively precise. The feedback was given based on the proportional error = |t_r_ − t_s_ |/t_s_, where t_s_ is the sample duration and t_r_ is the reproduced duration. The criterion of the binary feedback was self-adapted for each participant so that each kind of feedback appeared in around 50% of trials. After presenting the feedback for 50 ms, the fixation point turned back to grey to indicate the start of a new trial.

Ten different sample durations formed a uniform distribution between 520 to 880 ms, with a minimal difference of 40 ms between every two sample durations. Different sample durations were pseudo-randomized within blocks of 100 trials. Each participant finished 20 blocks, with a minute rest between every two blocks and daybreak after the first 10 blocks, resulting in 2000 trials in total and 200 trials for each sample duration.

#### Data analysis

The reproduced durations shorter than 0.3 s or longer than 1.5s (less than 1 ‰) were seen as outliers and excluded from the analysis. In all experiments, the first section was seen as a familiarization phase and removed from further analysis. The individual’s averaged reproduced durations were calculated for different sample durations, respectively. The regression-to-mean is quantified as the regression coefficient between the reproduced durations and the sample durations. We applied an index named “deviation” to examine serial dependence. The individual’s averaged reproduced durations were calculated for different sample durations, respectively. The deviation was calculated by subtracting the individual’s averaged reproduced durations of the corresponding sample duration from the reproduced duration of each trial^32,33^. Positive and negative values indicate that the reproduced duration of the present trial was longer or shorter than the averaged reproduction, respectively.

#### Derivate of Gaussian (DoG)

To quantify the magnitude of the serial dependence, a simplified DoG curve was used to fit the relationship between the difference between the sample durations and the deviation:

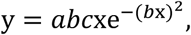

where y is the deviation, x is the relative sample duration of the previous trial, *a* is half the peak-to-trough amplitude of the derivative-of-Gaussian, *b* scales the width of the Gaussian derivative, and *c* is a constant 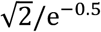, which scales the curve to make the *a* parameter equal to the peak amplitude. As a measure of serial dependence, we report half the peak-to-trough amplitude (half-height) and half the width of the best fitting derivative of a Gaussian. A positive value for the *a* parameter indicates a perceptual bias toward the sample duration of the previous trial. A negative value for the *a* parameter indicates a perceptual bias away from the sample duration of the previous trial. We fitted the Gaussian derivative at the group level and each individual using constrained nonlinear minimization of the residual sum of squares. Jackknife resampling was applied to estimate the variation of the parameters for the group-level fitting, where each participant was systematically left out from the collapsed sample. The standard deviation of those estimations represented the standard error of the parameter at the group level. To test the goodness-of-fit of our model, we computed ΔAIC of the DoG model compared with either a non-modal (y = 0, ΔAIC_n_) or a linear model (y = kx, ΔAIC_l_). A negative ΔAIC indicates DoG performed better than the alternative models.

To further analyze how many previous trials could influence the current reproduction, we fitted individual serial dependence of the 1-back (n-1), 2-back (n-2), and 3-back (n-3) trials with the DoG curve. Moreover, to check whether the deviation was a proper index to analyze serial dependence, we also tested the serial dependence of the 1-future (n+1) trial. Since individual serial dependence does not always show a strong non-linearity, *a* parameter (half-height) became unreasonably large in some cases. Therefore, we used the instantaneous slope of the DoG at the inflection point (*abc*), which is a suitable index to measure the sign of the serial dependence no matter whether participants showed a nonlinear curve or not.

### Experiment 2

#### Procedure and design

Experiment 2 was designed to test whether the serial dependence happened in the time of perception or motor response. To insert no-action trials where participants do not make a motor response, a cue should be presented between the end of the sample duration and the beginning of the reproduction to inform participants whether it was a no-action trial. Since this cue is easy to be confused with the “Ready” or “Set” signals and would interrupt the reproduction in the ready-set-go paradigm, we applied a different temporal reproduction task where participants perceived the presentation time of a stimulus but not the duration between two fleshes (“Ready”/ “Set”).

Each trial started with a fixation point at the center of the screen. After a random interval ranging from 0.7-1.2 s (following an exponential distribution), a ripple-shaped stimulus centering at the fixation point (see Fig.1c) was presented for sample duration and then disappeared. After 300 ms, the fixation point changed into either a “+”, that informed participant to reproduce the duration of the stimulus (action trial), or a “x”, which informed participants to make no response (no-action trial). In the action trials, participants could make a response at any time they want by holding the keypress for sample duration. After the key released, the fixation point changed color to present feedback for 300 ms in the same way as Experiment 1. In the no-action trial, the “x” presented for 700 ms. Then the fixation point changed back into a point indicating the start of a new trial.

Forty percent of trials of each sample duration were set as no-action trials, where participants did not perform reproduction after perceiving the sample duration. Instead, they viewed the fixation point for 700 ms. Five sample durations (540, 620, 700, 780, 860 ms) were used in Experiment 2 and formed a uniform distribution. A total of 1800 trials was included with 360 trials for each sample duration. The sample durations and action/no-action trials were pseudo-randomized within a block of 180 trials. Participants went through 5 blocks (900 trials) at the roughly same time each day, with a rest of one minute between every two blocks.

#### Data analysis

The reproduced duration shorter than 0.3 s or longer than 1.5 s (less than 1‰) were seen as outliers and ruled out from the further analysis. To analyze the serial dependence with and without motor response, the action trials were sorted into two kinds based on whether it was preceded by an action trial or a no-action trial. We calculated the average reproduced durations and the deviation using the same protocol as Experiment 1. We fitted DoG separately for two kinds of action trials to measure the serial dependence of the 1-back trial and calculated the instantaneous slope of the DoG at the inflection point. Note that if serial dependence observed in Experiment 1 was due to the motor response, we should expect no serial dependence in the action trial preceded by a no-action trial. Moreover, for the action trial preceded by a no-action trial, we also analyzed the serial dependence of the 2-back action trial. Group-level DoG fitting was performed on 1-back and 2-back action trials separately to quantify the amplitude of serial dependence.

### Experiment 3

#### Procedure and design

Experiment 3 was designed to further test whether serial dependence also happens in the perception even if no motor response is made. Experiment 3 used the same experimental paradigm (stimulus procedure) as Experiment 2. The sample duration of action trials was fixed at 700 ms. Zero to five no-action trials sampled from either the long or short prior conditions were inserted between every two action trials. The short prior condition contained 500, 550, 600, 650 ms. The long prior condition contained 750, 800, 850, 900 ms. All eight sample durations in short and long prior conditions appeared for equal times in the experiment. The probability of 0, 1, 2, 3, 4, 5 no-action trials between two action trials was 10%, 10%, 10%, 10%, 30%, 30%, respectively. Participants went through a total of 2064 trials, among which 480 trials were action trials. Prior conditions were pseudo-randomized within blocks of 516 trials (120 action trials), with a rest of one minute between every two blocks.

#### Data analysis

The reproduced durations shorter than 0.3 s or longer than 1.5s (less than 1‰) were seen as outliers and excluded from further analysis. Action trials were grouped based on the number of no-action trials ahead of them and whether those no-action trials belonged to the long prior or the short prior conditions. Action trials after 1, 2, and 3 no-action trials were grouped together to make each no-action trial number condition (1 to 3, 4, 5 no-action trials) contain 30% of action trials. The difference between the reproduced durations in long and short prior conditions was calculated for different numbers of no-action trials, respectively. One-sample t-tests were applied to compare the differences from zero.

### Experiment 4

#### Stimulus and design

We tested whether the range of stimuli influenced serial dependence. The stimulus and procedure of Experiment 4 were similar to Experiment 2, except that the fixation point always changed into “+”, and participants were told they should make a response to every trial. The sample durations of the median-range condition were 540, 670, 900, 1030, 1160 ms, with 150 trials for each sample duration, and the sample durations of the long-range condition were 560, 880, 1300, 1620, 1840 ms, with 120 trials for each sample duration. Different sample durations were pseudo-randomized within each of the five blocks, resulting in a total of 750 trials for the median-range and 600 trials for the long-range conditions. With these trial numbers, both two experiments lasted for around one hour.

#### Data analysis

The outliers of the reproduced durations were set as those shorter than 0.3 s or longer than 2.5s (less than 1 ‰) and were excluded from further analysis. The half-height and half-width of the best-fitted DoG were compared between groups with t-test, where the t-value was computed with the mean and the variance estimated from the jackknife resampling. To compare serial dependence between different duration ranges, t-tests were applied for median-range verse Experiment 1, median-range verse Experiment 2, and median-range verse long-range. Bonferroni correction was applied for multiple comparisons. Moreover, to examine how the lengths of stimulus durations influence serial dependence within one range, we grouped the stimulus durations into pairs with the same distance and analyzed the influence of the 1-back trial when both the current and the 1-back stimulus durations belonged to the pair. For example, the pair of 560 & 880 ms includes trial in two scenarios: 1) trial n-1 was 560 ms and trial n was 880 ms; 2) trial n was 560 ms and trial n-1 was 880 ms. The averaged deviation of the short stimulus in the pair (scenarios 2) was subtracted by the averaged deviation of the long stimulus (scenarios 1) and then divided by two to calculate the magnitude for serial dependence (paired half-height). The median-range was grouped into three stimuli pairs (540 & 900 ms, 720 & 1080 ms, 900 & 1260 ms) since the distance in each pair (360 ms) was close to twice the half-width. For long-range, every two successive stimuli were tied into one pair (560 & 880, 880 & 1200, 1200 & 1520, 1520 & 1840 ms). Repeated ANOVAs were applied to compare the paired half-heights among different stimulus pairs.

### Experiment 5

#### Stimulus and design

To further examine how the variability of stimuli durations influences serial dependence, two new contexts were included to compare with the median-range in Experiment 4: 1) A new short-range condition with stimuli durations ranging from 720 to 1080 ms with a step of 40 ms; 2) A new median-range condition with stimuli following a bimodal distribution. This new short-range condition shared the same mean as the median-range condition and shared the same range as Experiment 1. Trials were pseudo-randomized within each of the ten 100-trial blocks, resulting in a total of 1000 trials. The bimodal-distribution condition shared the same stimuli durations as the median-range in Experiment 4 (560, 880, 1300, 1620, 1840 ms). However, the trial number of 560 and 1840 ms was three times of the others (880, 1300, 1620 ms), which induced a larger contextual variability than the uniform distribution in the previous median-range condition. Trials were pseudo-randomized within each of the ten 135-trial blocks, resulting in a total of 1350 trials. All analyses followed Experiment 4.

### Models

#### Adaptation-level (AL) model

The AL model assumes that the internal reference in temporal perception is represented as a single vale (t_o_). In each trial, the estimated duration (t_e_) was a weighted summation of t_o_ and the sample duration (t_s_).

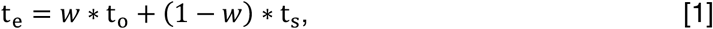

where the weight *w* represents how much participants rely on the reference point. A larger *w* caused a stronger central tendency. The observers make a motor response (t_r_) based on t_e_.

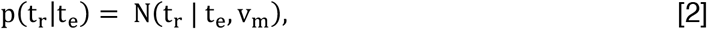

where N(x|m, s) represents a normal distribution with mean m and standard deviation s. t_r_ is the reproduced duration, and v_m_ scales the motor noise. After the motor response, t_0_ is updated based on t_r_ linearly:

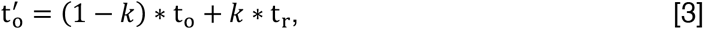

where 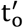 is the new reference point for the next trial, and *k* represents the learning rate.

#### Bayesian Recency (BR) model

The BR model assumes that the internal reference is represented as a prior distribution, π(t_s_), constructed from the sample durations. The likelihood function describes the probability relation between the perceived duration (t_p_) and t_s_.

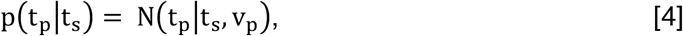

where v_p_ scales the perception noise. The posterior, π(t_s_|t_p_), is the product of the prior multiplied by the likelihood function and appropriately normalized.

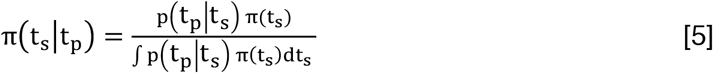

The loss function, l(t_e_, t_s_), was used to convert the posterior into a single estimation, t_e_, which is the mean of the posterior in our case.

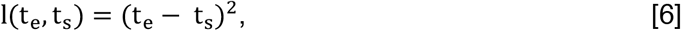

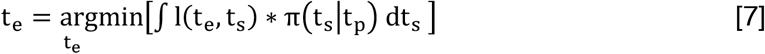

Similar to the AL model, the Bayesian observer makes motor responses based on t_e_ :

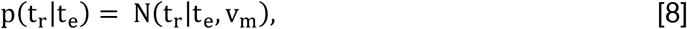

where t_r_ is the reproduced duration, and v_m_ scales the motor noise. Since previous studies^11,16^ indicated that the scalar form of perception noise and motor noise fitted participants’ behavior better than the models setting v_p_ and v_m_ as constants, we set v_p_ as n_p_ * t_s_, and v_m_ = n_m_ * t_e_, where n_p_ and n_m_ are constants. These assumptions were consistent with the theory of scalar property in interval timing^11,31^.

After each trial, the internal reference is updated according to t_r_. Following the theory of predictive coding, the observer assumes that the sample duration is more likely to appear around t_r_ in the future trials, so they add a scaled normal distribution centered at t_r_ to the prior and then normalized:

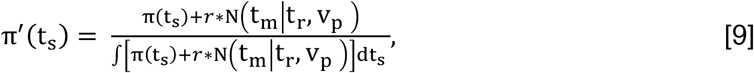

where π′(t_s_) is the new prior distribution (internal reference), and *r* is the learning rate.

#### Bayes-Least-square (BLS) model

This model was taken from the Bayesian observer model^16^ with Bayes-Least-square as the mapping rule. The procedures to infer t_m_ from t_s_ and π(t_s_) were the same as the BR model mentioned above (formula [4]-[7]). The only difference was that the true distribution of sample durations was applied as the internal reference:

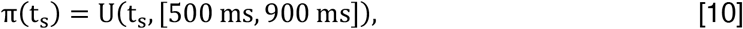

where U(x, [y, z]) represents a uniform distribution ranging from y to z. The observers are assumed to always apply this uniform distribution as prior and never updated it.

#### Bayesian Integration (BI) model

The BI model was adapted from the previous study on the sequential effect in numerous perception^30^. This model suggests that the observer assigned weights between the previous response (t_r_) and the current stimulus (t_s_) when estimating the current duration:

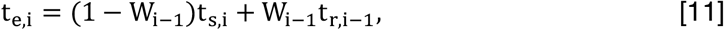

where t_e,i_ and t_s,i_ indicate the t_e_ and t_s_ of the trial i, and t_r,i−1_ is the reproduced duration of trial i-1. W_i−1_ is the weight the observer assigned to the 1-back response when estimating the current duration. Following the Bayesian rule to integrate two observations of the same stimulus, the observer will specify W_i−1_ as

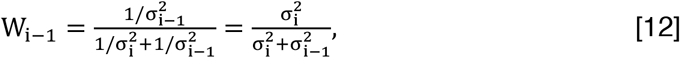

where σ is the variance of the estimation. However, two continuous stimuli are not identical in our case, so the weight is influenced by the distance between two stimuli,

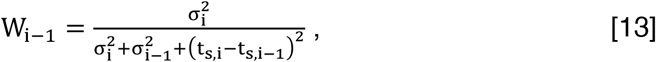

The variance is supposed to follow a power law.

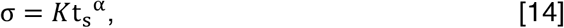

Similar to the BR model, we suppose temporal perception follows the scaled property in our experiments, then *α* = 1. *K* is the free parameter regulating the behavior of the model.

#### Heretical Bayesian Integration (HBI) model

The HBI is a combination of the BLS and BI. It contained two Bayesian processes. In the first Bayesian integration, the observers integrate the current stimulus (t_s_) with prior π(t_s_) to get an estimation t_e_′, following the formula [4]-[7]. The second Bayesian integration computes t_e,i_ based on t_r,i−1_, t_e,i_′ and t_e,i−1_′ following the rules in the BI model. The formulas [11] and [13] are rewritten as

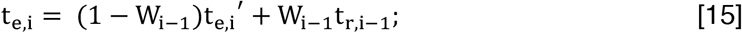

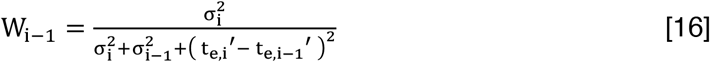

#### Simulation procedure

All models were applied to simulate the result of Experiment 1. 100 pseudo-participants were generated to perform the task in each simulation. To determine values of the free parameters for model simulation, we referred to a previous study that fitted the BLS in a design very similar to our experiment^16^. Based on their results, we set n_p_= 0.10 and n_m_ = 0.06 for the BLS, the BR, and the HBI. Other parameters were determined to make the amplitude of serial dependence roughly similar to the behavioral results. Specifically, in the AL model, we set *w* = 0.7 and *k* = 0.3. In the BR, the learning rate *r* was specified to be 0.3. In the BI and the HBI, *K* was set as 0.06. To further compare between BR and HBI, two models both showing central tendency and non-linear serial dependence, we simulated the prediction of them for different stimulus duration contexts. For simulations of short-range, the step between each two adjacent stimuli durations was 40 ms. For simulations of the median-range and the long-range conditions, to gain a higher resolution, the step of stimuli was set as 70 ms instead of the actual steps applied in behavioral experiments. The n_p_ and n_m_ were kept the same as the previous simulation, while a series of *r* and *K* was applied. For the illustration purpose, in Fig. 6d&n, a large learning rate r = 0.7 was used to show a more obvious difference between conditions.

## Acknowledgments

This work was supported by the National Natural Science Foundation of China (31771213 and 31371018) to Yan Bao. We thank Jiashu Wang, Huihui Zhang, and Hang Zhang for valuable discussions.

## Author contribution

T.W. and Y.B. designed the experiment. T.W. and Y.L. collected the data. T.W. analyzed the data. T.W. prepared the figures. T.W., E.P, and Y.B. involved in writing the manuscript with the initial draft written by T.W.

